# A PLK1-Directed Mitotic Genome Maintenance Network Promotes DNA Synthesis in Mitosis

**DOI:** 10.64898/2026.07.06.736709

**Authors:** Daniel-Cristian Ungureanu, Ivan Muñoz, Wissem Bououdina, Szymon A. Barwacz, Thomas Macartney, Frederic Lamoliatte, Yixin Wang, Ying Liu, Florian Weiland, John Rouse

**Affiliations:** MRC Protein Phosphorylation and Ubiquitylation Unit, University of Dundee, DD1 5EH; Division of Genome Stability, Faculty of Life Sciences, James Black Centre, University of Dundee, DD1 5EH; Centre for Chromosome Stability, Dept of Cellular and Molecular Medicine, University of Copenhagen, Copenhagen, Denmark; Department M2S, Centre for Food and Microbial Technology (CLMT), Laboratory of Enzyme, Fermentation and Brewing Technology, KU Leuven-Gent, 9000 Gent, Belgium

## Abstract

Polo-like kinase 1 (PLK1) is a master regulator of mitosis and is known to dictate DNA repair pathway choice at this stage of the cell cycle. However, its roles in controlling mitotic DNA damage responses remain incompletely characterised. Here, we used acute PLK1 inhibition as a substrate-trapping strategy to stabilise PLK1–target interactions in mitotic cells and identify PLK1-associated DNA repair factors. Proteomic analysis of endogenous HA-tagged PLK1 complexes revealed interactions with multiple genome stability proteins, including SLX4, RAD52, FANCM, and REV1. We demonstrate that PLK1 binds SLX4 and RAD52 via canonical CDK1-dependent phospho-docking motifs centred on SLX4 Ser1453 and RAD52 Thr300. Mutation of these residues abolished PLK1 binding and, at least for RAD52, prevented PLK1-dependent phosphorylation of mitotic targets. Functional analyses showed that PLK1 docking to SLX4 is dispensable for interstrand crosslink repair but essential for mitotic DNA synthesis (MiDAS), defining a separation-of-function allele. Likewise, disruption of PLK1 docking to RAD52 impaired MiDAS. Together, these findings identify PLK1 as a key coordinator of mitotic genome maintenance pathways required for MiDAS.

## Introduction

Polo-like kinase 1 (PLK1) is an essential serine/threonine protein kinase that controls entry to, and progression through, multiple stages of the cell division cycle in mammalian cells (Chapagai *et al*, 2025; Kalous & Aleshkina, 2023; Kim, 2022; Strebhardt, 2010). It plays a central role in regulating entry to mitosis, for example, by phosphorylating and activating key proteins involved in the G_2_/M transition. Once cells have entered mitosis, PLK1 is essential for proper centrosome maturation and bipolar spindle formation, and ensures accurate chromosome alignment and segregation by coordinating kinetochore-microtubule attachments. Furthermore, it regulates the spindle assembly checkpoint (SAC) and participates in cytokinesis by controlling contractile ring formation and abscission (Barr *et al*, 2004; Lane & Nigg, 1996; Petronczki *et al*, 2008; Sumara *et al*, 2004; Zitouni *et al*, 2014).

The phosphorylation of PLK1 substrates requires docking to its targets, which is mediated by its C-terminal polo-box domain (PBD) (Elia *et al*, 2003a; Elia *et al*, 2003b). The PBD functions as a phospho-recognition module that binds short motifs generated by prior phosphorylation events, most commonly catalysed by the major mitotic cyclin-dependent kinase, CDK1-cyclin B, at mitotic entry. Classical models describe this as a “priming” mechanism in which CDK1-dependent phosphorylation creates a consensus PBD-binding motif, enabling PLK1 recruitment and subsequent phosphorylation of the same substrate. Following PBD-mediated docking, PLK1 is positioned to phosphorylate nearby residues via its kinase domain, thereby enabling precise temporal coordination of mitotic processes (Elia *et al*., 2003a; Elia *et al*., 2003b; Esposito-Verza *et al*, 2026).

Beyond mitosis, it has become apparent that PLK1 regulates cellular responses to DNA damage predominantly, but not exclusively, in mitosis (Hyun *et al*, 2014). Polo-like kinase 1 (PLK1) plays a central role in coordinating cellular responses to DNA damage with mitotic progression, acting as both a driver of DNA damage checkpoint recovery and as a regulator of repair pathway choice in mitosis. Following genome damage in G_2_, checkpoint pathways - primarily mediated by ATR-CHK1 signalling - halt cell cycle progression to allow DNA repair. As cells recover, PLK1 becomes progressively reactivated and helps to switch off the checkpoint (van Vugt *et al*, 2004). To this end, PLK1 phosphorylates and targets key checkpoint components such as Claspin and CHK1 for inactivation or degradation, thereby dismantling the signalling apparatus required to sustain arrest of cells in G_2_ (Mamely *et al*, 2006; Peschiaroli *et al*, 2006). In parallel, PLK1 reinforces CDK1 activation, creating a positive feedback loop that drives irreversible mitotic entry even in the presence of residual DNA lesions (Gheghiani *et al*, 2017; Lindqvist *et al*, 2009).

Once cells have entered mitosis, the structural and temporal constraints of this phase seem to necessitate a fundamental rewiring of the DNA damage response. Chromosomes are highly condensed, potentially restricting access to repair machinery, while the rapid progression of mitosis might limit the time available for complex, high-fidelity pathways. In addition, the physical separation of sister chromatids reduces, in principle, the availability of homologous templates, which may render homologous recombination (HR) - a major pathway for the repair of double-strand breaks (DSB) - largely ineffective. Consequently, cells shift away from accurate but slower DSB repair mechanisms toward faster, more permissive pathways such as microhomology-mediated end joining (MMEJ) (Brambati *et al*, 2023; Giunta *et al*, 2010), which can operate on condensed chromatin without requiring extensive homology. Although error-prone, such pathways enable the rapid resolution of DNA lesions, highlighting how the unique features of mitosis drive a reprogramming of damage responses to prioritize completion of cell division over genomic fidelity.

A related manifestation of this adaptation in mitosis is mitotic DNA synthesis (MiDAS), a damage-tolerance pathway that allows cells entering mitosis with under-replicated genomic regions, or unresolved recombination intermediates, to complete DNA synthesis after bulk genome duplication has finished. MiDAS is particularly evident at difficult-to-replicate loci, including common fragile sites, and is stimulated by replication stress. Unlike canonical S-phase replication, MiDAS occurs on condensed mitotic chromosomes and depends on DNA repair and replication-associated factors such as MUS81-EME1, SLX4, RAD51, RAD52, PIF1, RTEL1, POLD3, REV1 and REV3 (Bhowmick *et al*, 2023; Minocherhomji *et al*, 2015). More recently it was shown that DNA polymerase theta (POLθ; POLQ) contributes to MiDAS via a microhomology-mediated break-induced replication (MM-BIR) pathway (Ngo *et al*, 2026). Thus, MiDAS provides a useful example of how cells repurpose genome maintenance factors during mitosis to process persistent replication intermediates before chromosome segregation is complete.

How MiDAS and other mitotic DNA damage responses are coordinated with cell-cycle progression remains incompletely understood, but Polo-like kinase 1 (PLK1) has emerged as a key candidate regulator of this mitotic rewiring (Hyun *et al*., 2014). PLK1 contributes the reported mitotic suppression of HR through the phosphorylation of CtIP, which prevents the interaction of this nuclease with DNA2, thereby suppressing long-range resection (Ceppi *et al*, 2023). PLK1 also inhibits the canonical non-homologous end joining (NHEJ) mode of DSB repair through phosphorylation of the XRCC4 subunit of DNA ligase IV (Terasawa *et al*, 2014). In addition to mitotic suppression of DNA repair pathways that dominate in interphase, PLK1 simultaneously promotes MMEJ. For instance, PLK1 regulates POLQ by phosphorylation events that enhance its recruitment to DSBs and favour this rapid, error-prone repair mechanism in mitosis (Gelot *et al*, 2023). PLK1 also phosphorylates RHINO, a component of the 9-1-1 complex, which promotes RHINO interaction with POLQ, and recruitment of POLQ to DSB sites which favours MMEJ (Brambati *et al*., 2023). Together, these activities position PLK1 as a key molecular switch that actively reshapes the DNA repair landscape in mitosis. There are hints that PLK1 may regulate DNA repair in mitosis by mechanisms that remain to be understood. For example, the scaffold protein SLX4, which coordinates three different structure-selective DNA repair endonucleases, interacts with PLK1 in mitosis but the impact of PLK1 docking on SLX4 function is not yet clear (Fekairi *et al*, 2009; Guervilly & Gaillard, 2018; Munoz *et al*, 2009; Svendsen *et al*, 2009).

In this study we took advantage of our observation that PLK1 inhibitors greatly increase the association of SLX4 with PLK1, which raised the possibility that these conditions “glue” PLK1 to its targets because kinase activity is required for substrate turnover. Here we purified endogenous PLK1 from mitotic cells treated with PLK1 inhibitors. Mining the resulting PLK1 interactors for GO terms connected with DNA damage responses identified a range of DNA repair proteins that could be subject to PLK1-mediated control in mitosis. We characterize two proteins that dock PLK1 in a CDK1-dependent manner in mitosis – SLX4 and RAD52 – and characterize these interactions and define SLX4 and RAD52 mutations that abolish interaction with PLK1. We show that the PLK1 docking on both SLX4 and RAD52 is essential for mitotic DNA synthesis (MiDAS).

## Results

### A strategy for identifying DDR targets of PLK1 in mitosis

We set out to identify the full range of DNA damage response proteins that are targeted by PLK1. To this end we took advantage of an observation we made concerning the PLK1-SLX4 interaction, in experiments where we co-immunoprecipitated SLX4 from extracts of cells arrested in prometaphase using either nocodazole (Zieve *et al*, 1980) or S-trityl-L-cysteine (STLC) (Skoufias *et al*, 2006), followed by a brief treatment with the highly selective PLK1 inhibitors BI-2536 or BI-6727 (Rudolph *et al*, 2009; Steegmaier *et al*, 2007). Antibodies recognizing the phosphorylated form of the microtubule-stabilizing protein TCTP (pSer46), a well-documented PLK1 target, confirmed PLK1 inhibition under the conditions we used (Cucchi *et al*, 2010). We noticed that exposure to PLK1 inhibitors led to a marked increase in the SLX4–PLK1 interaction (Fig. 1A), which suggested to us that inhibition of PLK1 kinase activity might stabilize its association with substrates by preventing phosphorylation-dependent dissociation, as suggested previously (Kishi *et al*, 2009). Building on this insight, we immunoprecipitated PLK1 from mitotic cells treated with PLK1 inhibitors, aiming to identify a broader set of PLK1-interacting proteins which could be mined for DNA repair-related factors.

**Figure 1.**
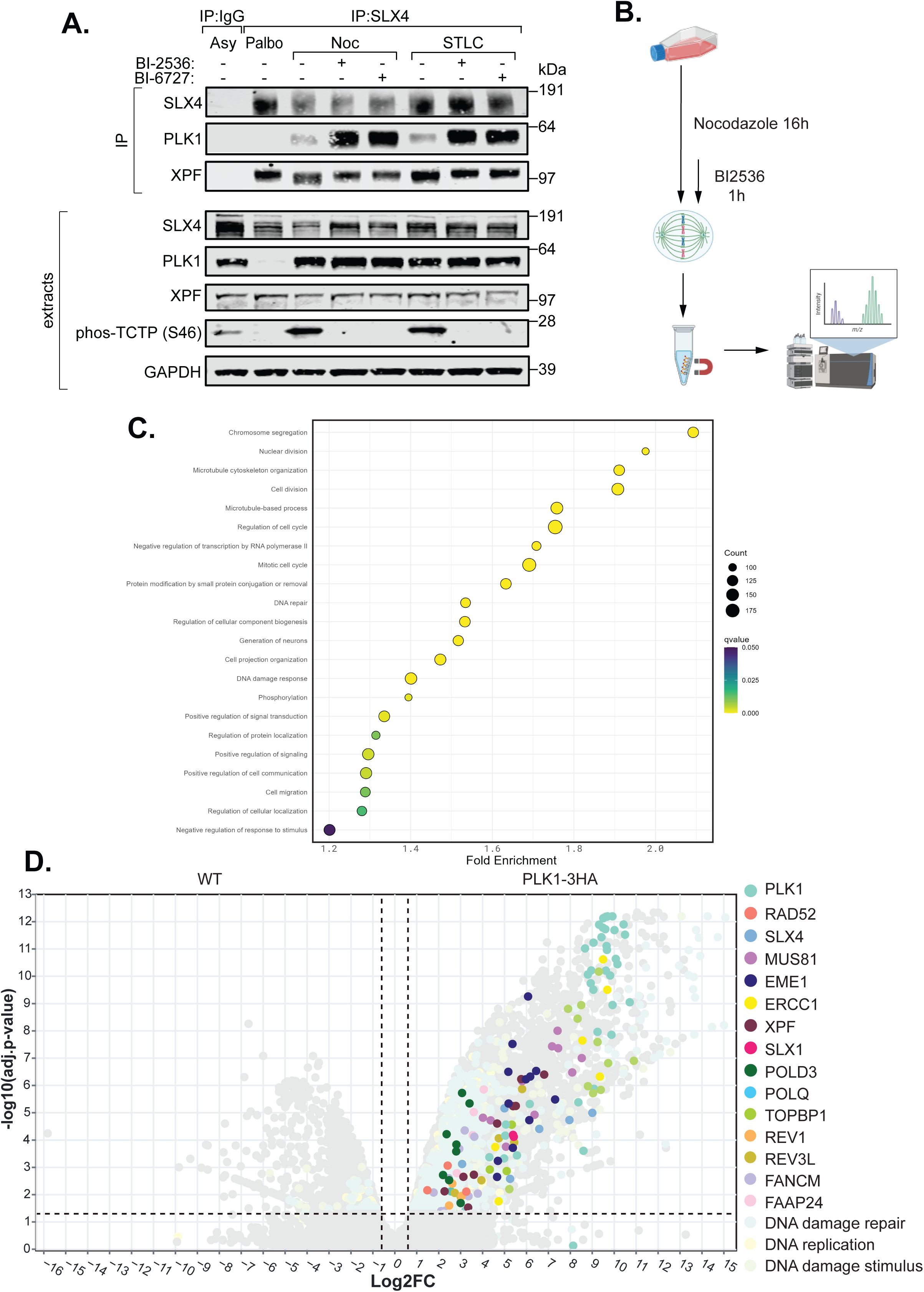
Identifying PLK1 binding proteins during mitosis. **A.** Extracts from U-2 OS cells that were either asynchronous or arrested with palbociclib (1 µM, 20h), nocodazole (150 ng/ml, 20h), or STLC (5 µM, 20h) were incubated with DMSO, BI-2536 (500 nM for 1h), or BI-6727 (500 nM for 1h), and subjected to SLX4 immunoprecipitation. Immunoprecipitates and whole-cell extracts were analysed by immunoblotting with the indicated antibodies. **B.** HCT116 parental and PLK1-3HA cells were treated with nocodazole (150 ng/ml) for 16h. In the last hour of nocodazole treatment, the cells were also treated with BI-2536 (500 nM) for 1h. The cell extracts were then subjected to HA immunoprecipitation coupled with mass spectrometry. **C.** Gene ontology (GO)-term enrichment analysis of the mitotic PLK1-3HA interactome showing log_10_ fold enrichment of the indicated GO terms in the PLK1-3HA cells relative to the parental cells. **D.** Volcano plot of mass spectrometry analysis of HA immunoprecipitates from 6 biological replicates, showing peptides differentially enriched in HA IPs from parental (WT) cells (left) or PLK1-3HA cells (right). The horizontal cutoff line represents a P-value of 0.05. Positive log2FC indicates higher abundance of the respective peptides in PLK1-3HA (vs parental untagged cells, and vice versa). Horizontal dashed line marks an adjusted p-value cut-off of 0.05 (5% FDR). Vertical lines correspond to a fold-change of (±) 1.5. Peptides situated on or outside these lines are considered differentially abundant. Colored dots indicate affiliation to the respective GO terms or proteins of interest as shown.

To facilitate isolation of PLK1, we employed CRISPR/Cas9 to insert a DNA sequence encoding a C-terminal 3XHA epitope tag into the *PLK1* locus in human HCT-116 cells. As illustrated in Figure S1A, we successfully obtained several clones with a single PLK1 allele tagged with HA, although, even after screening nearly 200 colonies, fully homozygous knock-in clones could not be recovered. Efforts to tag PLK1 at the N-terminus produced similar results (data not shown). Nonetheless, SLX4 was readily detected in anti-HA immunoprecipitates from mitotic extracts of cells expressing 3HA-tagged PLK1 (Figure S1B). Based on these findings, we proceeded with large-scale immunoprecipitation of PLK1-3HA from mitotic cells treated with BI-2536, employing six biological replicates, followed by mass spectrometry to identify candidate PLK1 substrates and interacting partners (Figs. 1B, S1C).

Our proteomic analyses identified 1,005 proteins enriched in PLK1-3HA precipitates (5% FDR) (Table S1), along with 168 proteins that were detected exclusively in PLK1-3HA samples, totalling 1,173 proteins. Gene ontology analysis yielded 447 significantly enriched terms (5% FDR). We grouped these into 22 clusters based on semantic similarity (Wang *et al*, 2007b; Yu *et al*, 2010), filtering for terms associated with more than 90 proteins (Fig. 1C). The GO terms most strongly represented among the PLK1 targets we identified are those relevant to the chromosome segregation (GO0007059), mitotic cell cycle (GO0000278) and microtubule-based processes (GO0007017) which is encouraging given the central role PLK1 plays in regulating these aspects of cell function. Notably, proteins linked to GO terms for DNA damage response (GO0006974), DNA replication (GO0006260), or DNA damage repair (GO0006281) were also represented including SLX4, POLQ and RHINO (Figs. 1C, 1D, 2A).

**Figure 2.**
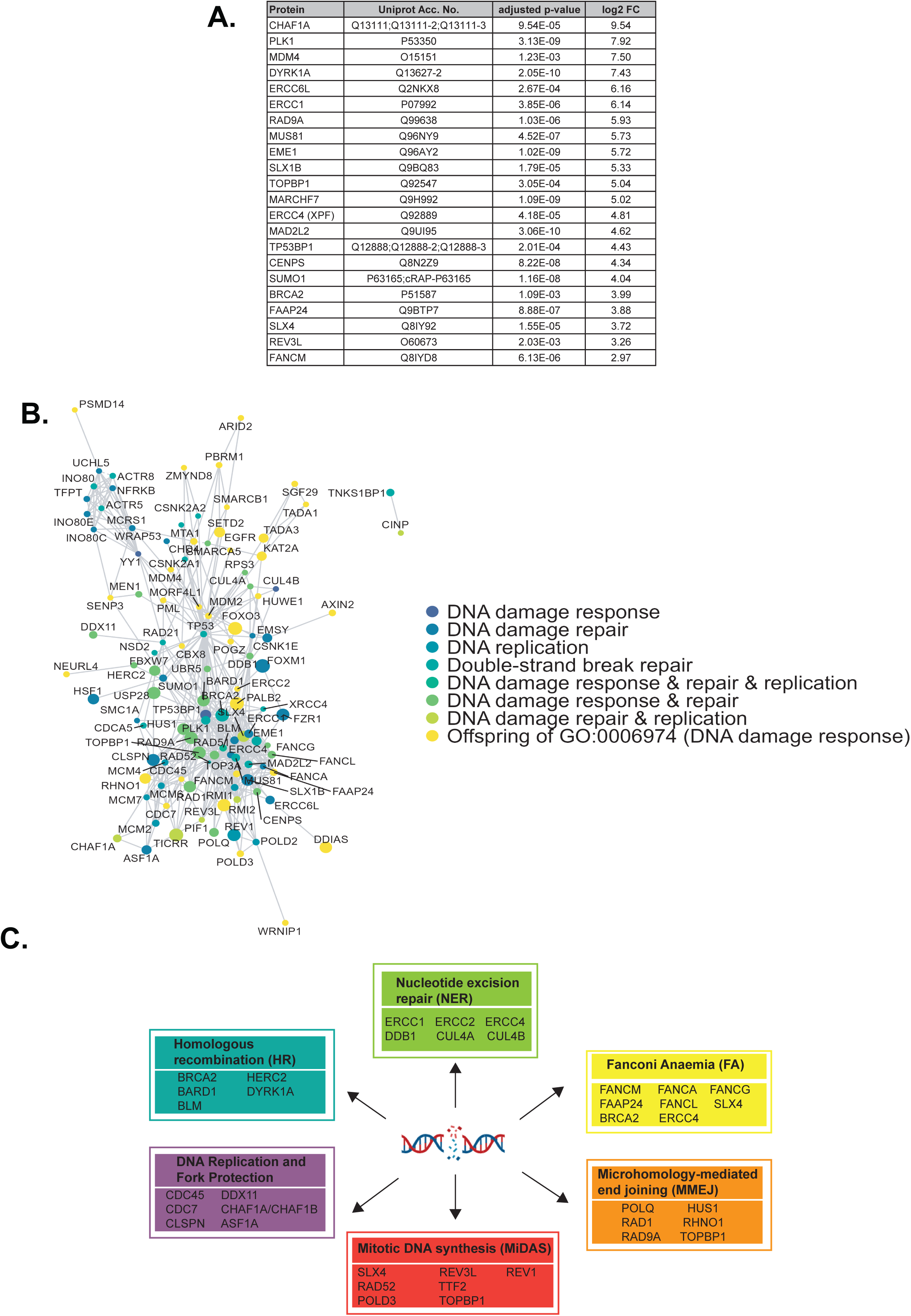
PLK1 interacts with a wide range of genome stability proteins in mitosis. **A.** List of the top twenty PLK1 interactors relevant to the GO terms listed in B below. **B.** Network plot of proteins higher in abundance in PLK1-3HA precipitates (adjusted p-value ≤ 0.05), including proteins which were detected in at least 4 out of 6 replicates which were not detected in precipitates from parental cells (all 6 replicates). The network was filtered to contain only proteins which exhibited at least one of the following GO terms: GO0006974 (DNA damage response), GO0006260 (DNA replication), GO0006281 (DNA damage repair) or their combinations. “Offspring of GO0006974” indicates that the protein is associated with a GO term which is an offspring of GO0006974. Node size corresponds to the difference in average VSN transformed intensity of the protein in PLK1-3HA precipitates and the protein in negative control precipitates (parental cells). In case of no detection of the protein in all 6 negative control replicates, the corresponding average VSN intensity was set to 0. Edges indicate known protein-protein interaction, as derived from the STRING database, exhibiting a combined score of at least 700 (“high confidence”). **C.** Grouping a selection of the DDR hits into DNA repair pathways.

### Validating new PLK1 targets in genome maintenance

In addition to the known PLK1 targets SLX4, POLQ and RHINO, genome stability regulators not previously linked to PLK1 were identified in our mass spectrometric analyses of PLK1-3HA precipitates, including the SLX4-associated nucleases MUS81-EME1 and XPF-ERCC1, as well as the FANCM DNA translocase and the TLS polymerase REV1 (Figs. 1C, 2A). This group formed a strong interaction network, with TP53 and an SLX4/EME1/ERCC1/MUS81/PLK1 subnetwork at the core (Fig. 2B). Many of the genome stability-related PLK1 targets identified in our screen fell into well-characterized genome maintenance pathways (Fig. 2C). Among these were the Fanconi anemia (FA) pathway, which is crucial for the repair of DNA interstrand crosslinks (ICLs) and replication fork stalling responses (Deans & West, 2011); nucleotide excision repair, responsible for removing bulky base lesions such as UV-induced DNA damage (Zhang *et al*, 2022); homologous recombination (HR), which repairs double-strand breaks and collapsed replication forks (Reginato & Cejka, 2020); and mitotic DNA synthesis (MiDAS), a pathway that addresses under-replicated genomic regions during mitosis (Bhowmick *et al*., 2023; Minocherhomji *et al*., 2015). We also found proteins involved in DNA replication and replication fork stability (Fig. 2C).

To validate selected protein–protein interactions, we turned to western blot analysis. As shown in Fig. 3A, several proteins were confirmed in PLK1-3HA precipitates, including the DNA translocase FANCM (remodels replication forks, repairs ICLs) (Abbouche *et al*, 2024), the DNA repair scaffold protein SLX4 (also known as FANCP; involved in ICL repair, fork processing, and MiDAS) (Guervilly & Gaillard, 2018), ERCC1 (the regulatory subunit of the ERCC1-XPF nuclease, functioning in NER and ICL repair), and EME1 (the regulatory subunit of the EME1-MUS81 nuclease, involved in fork processing and MiDAS) (Ciccia *et al*, 2008). We also detected RAD52, a protein implicated in MiDAS and double-strand break repair (Fig. 3A) (Rossi *et al*, 2021). The interaction between each protein and PLK1-3HA was substantially stronger in nocodazole-arrested cells compared to asynchronous cells and was abolished by the CDK1 inhibitor RO3306, consistent with these proteins being *bona fide* PLK1 targets (Fig. 3A).

**Figure 3.**
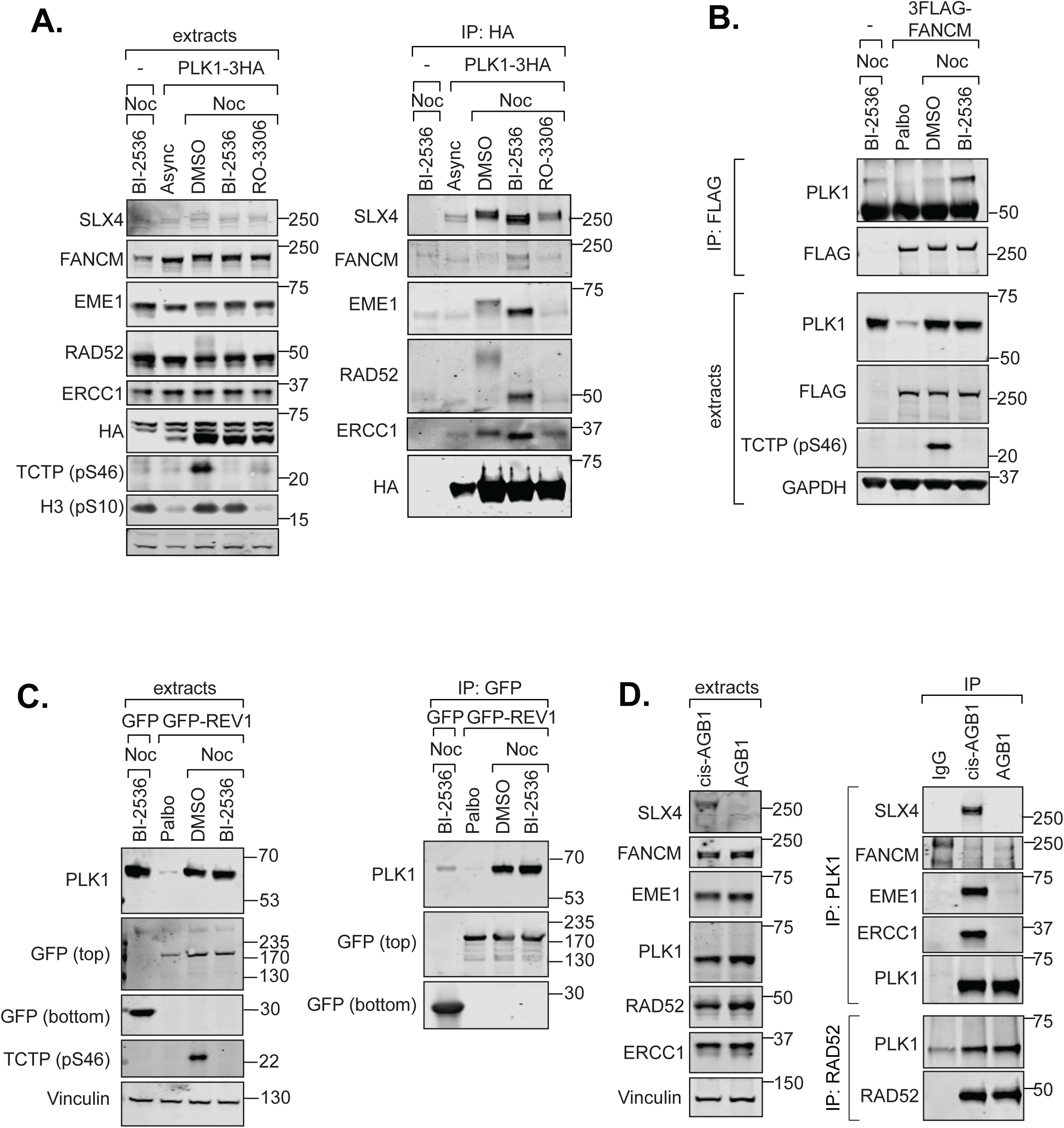
Validation of genome stability factors as mitotic PLK1 targets. **A.** Extracts from HCT116 parental and PLK1-3HA cells treated as described were subjected to HA immunoprecipitation and immunoblotted for the indicated proteins. **B.** U-2 OS FRT parental and 3FLAG-FANCM cells were treated with palbociclib (1 ìM) or nocodazole (150 ng/ml) for 20 hours. In the last hour of treatment, nocodazole-arrested cells were treated with DMSO or BI-2536 (500 nM) for 1 hour. Extracts from these cells were subjected to FLAG immunoprecipitation, followed by immunoblotting for the indicated proteins. **C.** U-2 OS FRT GFP and GFP-REV1 cells were treated as described in (B). Extracts from these cells were subjected to GFP immunoprecipitation, followed by immunoblotting for the indicated proteins. **D.** Extracts from nocodazole-arrested U-2 OS BromoTag-SLX4 cells treated with cis-AGB1 (300 nM, 1h) or AGB1 (300 nM, 1h) and BI-2536 (500 nM, 1h) were subjected to PLK1 and RAD52 immunoprecipitation and immunoblotted for the indicated proteins.

It is noteworthy that FANCM was identified in PLK1-3HA precipitates from mitotic cells only after a short incubation with BI-2536. This observation may reflect rapid, phosphorylation-dependent dissociation of the interaction under normal conditions. To investigate this further, we overexpressed a 3FLAG-tagged version of FANCM in U-2 OS cells. Consistent with our earlier findings, PLK1 was detected in 3FLAG-FANCM precipitates exclusively following BI-2536 treatment of nocodazole-arrested cells, providing additional support for FANCM as a PLK1 target (Fig. 3B). We also sought to validate the interaction between PLK1 and REV1, a specialized Y-family DNA polymerase that facilitates translesion synthesis and MiDAS (Sale, 2013; Wu *et al*, 2023). To this end, we expressed a GFP-tagged REV1 construct in U-2 OS cells. In this system, PLK1 was readily observed in GFP-REV1 precipitates from nocodazole-arrested cells, whereas only a weak signal was detected in cells arrested in G1 with palbociclib (Fig. 3C).

EME1-MUS81 and ERCC1-XPF nucleases are well-established interactors of SLX4 (Fekairi *et al*., 2009; Munoz *et al*., 2009; Svendsen & Harper, 2010). We wanted to determine if their association with PLK1 during mitosis was mediated indirectly via SLX4. To this end, we used CRISPR/Cas9 to engineer an inducible degron at the N-terminus of SLX4 (Fig. S2), as generating homozygous knockout clones for SLX4 proved unsuccessful in a range of cell lines (data not shown). The BromoTag, a modified Brd4 bromodomain, binds the PROTAC AGB1, which recruits the E3 ligase subunit VHL, enabling targeted degradation of SLX4 (Bond *et al*, 2021) (Figs. S2A-C). We isolated several homozygous clones with all SLX4 alleles BromoTagged, which showed complete SLX4 degradation after AGB1 treatment (Fig. S2D-F). Treatment with cis-AGB1, a derivative that does not bind VHL (Bond *et al*., 2021), had no effect on BromoTag-SLX4 levels. Complete degradation occurred within one hour of AGB1 treatment and was blocked by the 26S proteasome inhibitor MG132 (Rock *et al*, 1994) or MLN4924 (which inhibits cullin-RING ligases (Soucy *et al*, 2009), including SCF^VHL2^ (Figs. S2G, H). As illustrated in Fig. 3D, BromoTag-SLX4, EME1, and ERCC1 were robustly detected in PLK1 immunoprecipitates from nocodazole-arrested cells treated with BI-2536. However, following AGB1-mediated SLX4 degradation, EME1 and ERCC1 disappeared from PLK1 precipitates, as well as SLX4, indicating their association with PLK1 is dependent on SLX4. Notably, FANCM and RAD52 association with PLK1 was maintained even after SLX4 depletion (Fig. 3D).

As shown in Fig. 3, FANCM, EME1, ERCC1, SLX4, and RAD52 each exhibited electrophoretic mobility shifts in nocodazole-treated cells compared with asynchronous controls, with shifts varying in prominence. These shifts were reversed upon treatment with BI-2536. To further characterise these mobility changes, we ran samples on different types of PAGE gels. As illustrated in Fig. S3, the PLK1-dependent shifts in FANCM and SLX4 were most pronounced on Tris-acetate gels, whereas the mobility shifts for RAD52 and EME1 appeared consistent across all gel types tested. Taken together, the requirement for CDK1-dependent docking and the observation of PLK1-dependent phosphorylation shifts strongly support the conclusion that these proteins are directly targeted by PLK1.

Altogether, our findings identify several previously unrecognized genome stability regulators as mitotic targets of PLK1. Building on these results, we next explored in greater depth the regulation of two such proteins - SLX4 and RAD52 - by PLK1.

### PLK1 interacts with CDK1-phosphorylated SLX4 phospho-Ser1453 *in vitro* and in cells

The PLK1 polo-box domain (PBD) acts as a phospho-recognition module, binding short motifs in target proteins created by CDK1-dependent phosphorylation, classically Ser-pSer/Thr-Pro (Elia *et al*., 2003a; Elia *et al*., 2003b; Esposito-Verza *et al*., 2026). Human SLX4 contains four such motifs, centred on Ser960, Ser1244, Ser1355, and Ser1453 (Fig. 4A). To examine their role, we generated GFP-tagged SLX4 constructs with Ser-to-Ala substitutions at each site, testing their interaction with PLK1 in cells. As shown in Fig. 4B, the Ser1453Ala mutation abolished the interaction of GFP-SLX4 with endogenous PLK1, while mutations at Ser960, Ser1244, or Ser1355 had no apparent effect. This result pointed to a specific interaction between PLK1 PBD and SLX4 phosphorylated at Ser1453 by CDK1.

**Figure 4.**
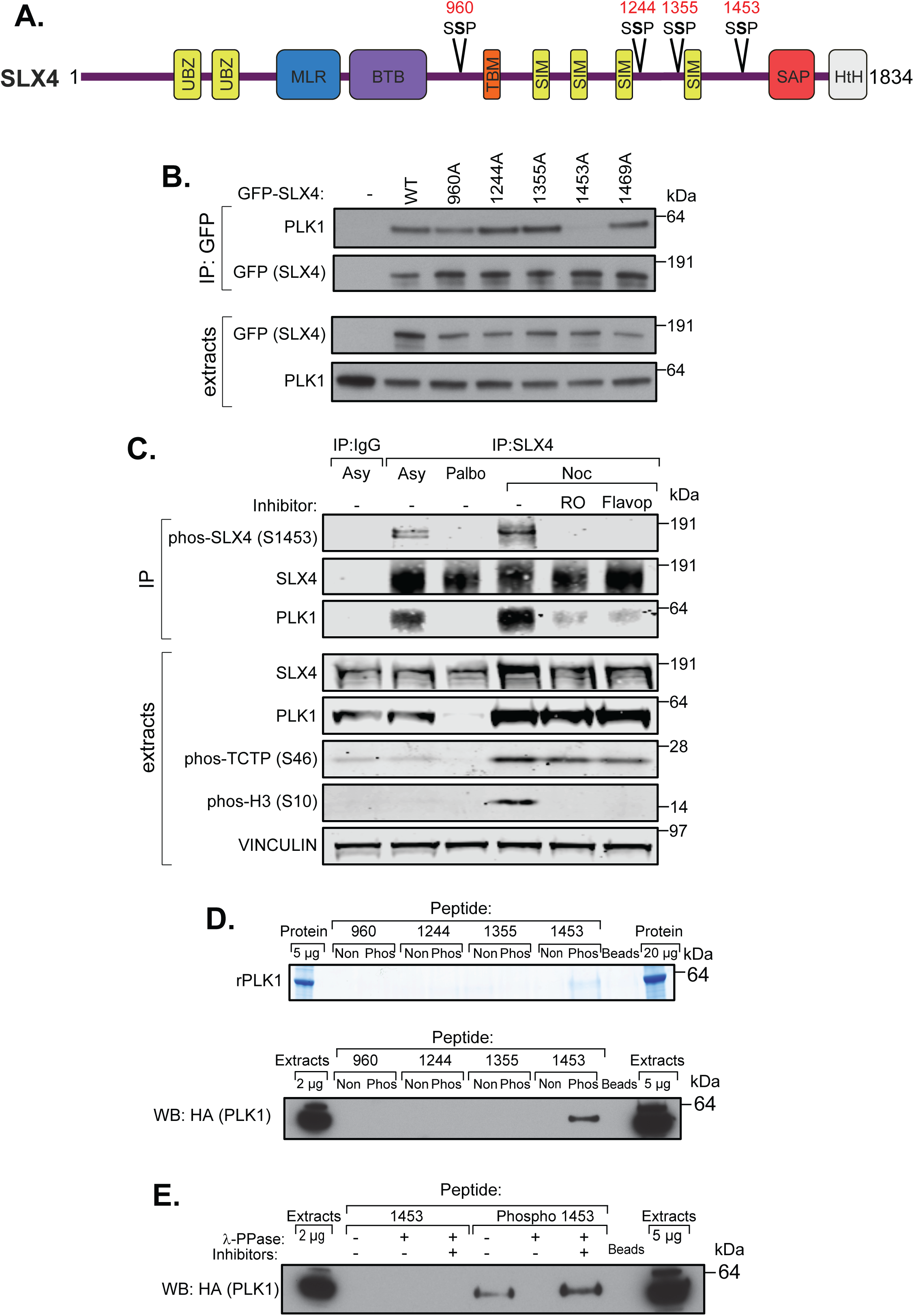
PLK1 binds directly to CDK1-phopshorylated Ser1453 in SLX4 in mitosis. **A.** Schematic representation of SLX4 showing conserved domains and candidate PLK1 PBD-binding motifs (S-(S/T)-P) **B.** Top panel: Bacterially expressed recombinant hexahistidine-tagged PLK1 (Munoz *et al*., 2018) was incubated for 2 h at room temperature with streptavidin beads coupled to the indicated peptides. Beads were washed extensively, and bound proteins were eluted in LDS sample buffer. Pull-downs were resolved by SDS-PAGE and visualized by Coomassie Brilliant Blue staining. Bottom panel: As in the top panel, except that streptavidin beads were incubated with extracts from HEK293 cells expressing PLK1-3HA, and precipitated proteins were analysed by immunoblotting with anti-HA antibodies. **C.** Extracts from HEK293 cells expressing PLK1-3HA were incubated with streptavidin beads coupled to the indicated peptides. Beads were washed extensively and resuspended in 1× lambda phosphatase buffer, followed by mock treatment or incubation at 37°C for 30 min with lambda phosphatase, or with lambda phosphatase pre-inactivated by treatment with microcystin and EDTA (“inhibitors”). Precipitates were resolved by SDS-PAGE and analyzed by immunoblotting with anti-HA antibodies. **D.** Schematic of human PLK1 showing domain organization, catalytic residue K82, and phospho-pincer residues H538 and K540 **E.** *Top panel*. Extracts from HEK293 cells expressing wild type PLK1-3HA or PLK1-3HA bearing indicated mutations were incubated with streptavidin beads coupled to the indicated peptides. Precipitates were analysed by immunoblotting with anti-HA antibodies. *Bottom panel*. Extracts from HEK293 cells expressing wild-type PLK1-3HA or PLK1-3HA bearing the indicated mutations were immunoprecipitated using anti-HA beads. Immunoprecipitates and whole-cell extracts were analyzed by immunoblotting with the indicated antibodies.

To further investigate, we developed sheep antibodies against phosphorylated SLX4 Ser1453, which proved highly selective for the phosphorylated immunogen peptide (Fig. S4A). Recognition of GFP-tagged SLX4 immunoprecipitated from nocodazole-arrested U-2 OS cells by this antibody was abolished by lambda phosphatase but not when a phosphatase inhibitor cocktail was present (Fig. S4B). As shown in Fig. S4C, GFP-tagged SLX4 isolated from G1-arrested (palbociclib-treated) cells reacted weakly with the pSer1453 antibody, but this signal increased dramatically after nocodazole arrest. Importantly, the Ser1453A SLX4 substitution prevented antibody recognition (Fig. S4C). Furthermore, the SLX4 pSer1453 signal in nocodazole-arrested cells was strongly reduced by CDK1 inhibitors RO3306 (Vassilev *et al*, 2006) or flavopiridol (Stadler *et al*, 2000), which also reduced SLX4 interaction with PLK1 (Fig. 4C). Together, these data establish that PLK1 interacts with SLX4 in mitosis via CDK1-dependent phosphorylation of S1453, and mutation of this residue abrogates the PLK1-SLX4 interaction.

To determine if the interaction between SLX4 pSer1453 and PLK1 is direct, we synthesized biotinylated peptides corresponding to regions around the four PLK1 candidate docking sites in SLX4, each with or without phosphorylation at the relevant serine. Immobilized peptides were incubated with recombinant full-length human hexahistidine-tagged PLK1 produced in bacteria. As shown in Fig. 4D (top), PLK1 specifically bound the phospho-Ser1453 peptide, but not any of the other phosphopeptides or the non-phosphorylated S1453 peptide. Similarly, the SLX4 pSer1453 peptide, but not other phosphopeptides, was able to pull down SLX4 from cell extracts (Fig. 4D, bottom). Furthermore, treating the SLX4 phospho-Ser1453 peptide with lambda phosphatase abolished its ability to retrieve SLX4 from extracts, unless a phosphatase inhibitor cocktail was included (Fig. 4E).

Structurally, the PLK1 PBD comprises two tandem polo-boxes, PB1 and PB2, forming a single functional domain. PB2 is mainly responsible for recognizing phosphorylated docking motifs, with key residues His538 and Lys540 in human PLK1 contributing to a positively charged binding pocket that coordinates the phosphate group (Fig. 5A). These residues establish a network of hydrogen bonds and electrostatic interactions that stabilize the phospho-dependent interaction (Elia *et al*., 2003b). As shown in Fig. 5B, combined mutation of His538 and Lys540 abolished binding to the SLX4 phospho-Ser1453 peptide, while mutation of Lys82, which abolishes kinase activity (Lane & Nigg, 1996), had no effect. Moreover, His538Ala and Lys540Met mutations in PLK1 markedly reduced its interaction with SLX4 in mitotic cells (Fig. 5C). Taken together, these data demonstrate that the PLK1 PBD specifically recognizes a single CDK1 phosphorylation site, Ser1453, in SLX4 during mitosis, and this interaction is essential for the regulated association of SLX4 and PLK1.

**Figure 5.**
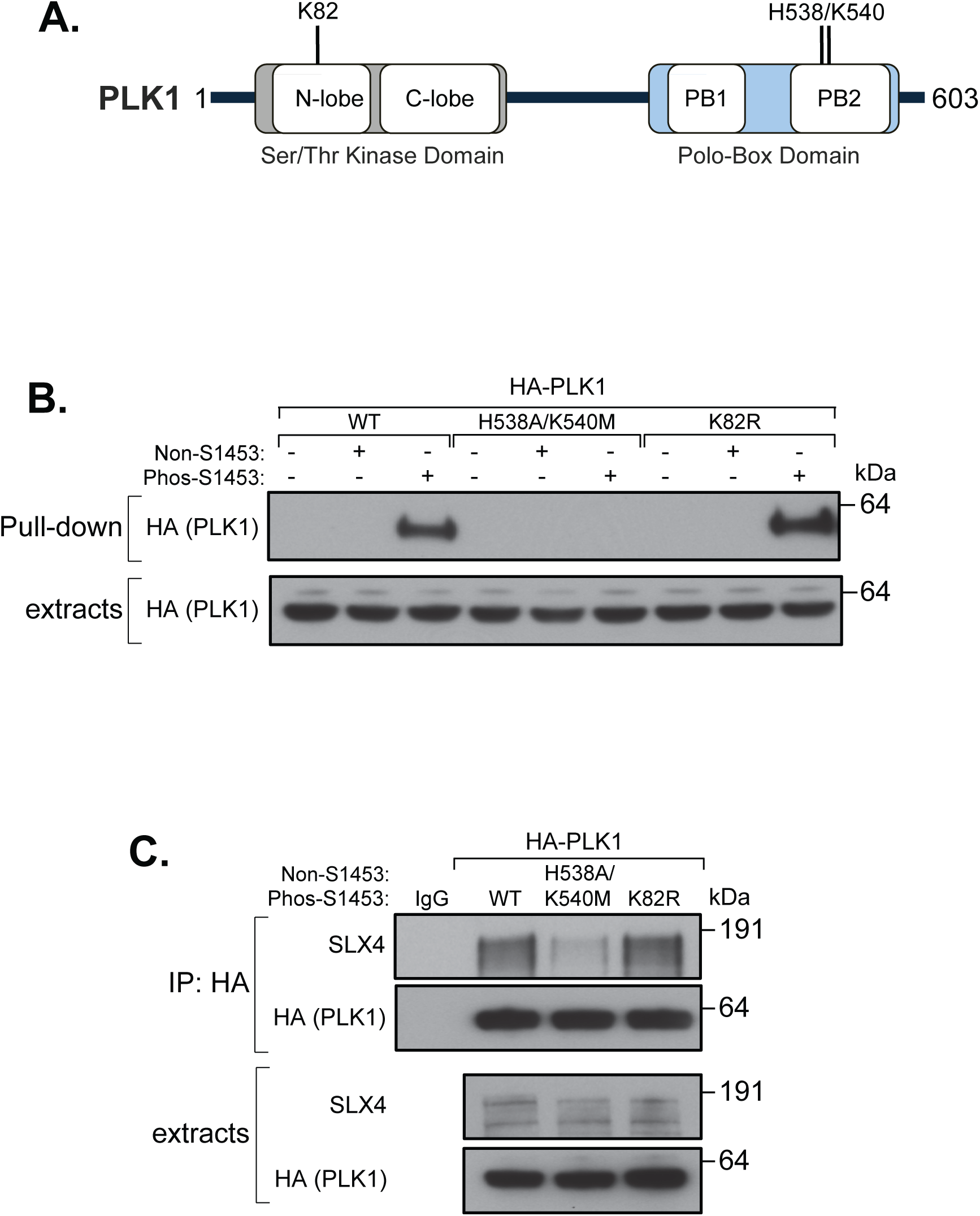
PLK1 polobox mutations abolish interaction with SLX4 pSer1453. **A.** Schematic of human PLK1 showing domain organization, catalytic residue K82, and phospho-pincer residues H538 and K540 **B.** Extracts from HEK293 cells expressing wild type PLK1-3HA or PLK1-3HA bearing indicated mutations were incubated with streptavidin beads coupled to the indicated peptides. Precipitates were analysed by immunoblotting with anti-HA antibodies. **C.** Extracts from HEK293 cells expressing wild-type PLK1-3HA or PLK1-3HA bearing the indicated mutations were immunoprecipitated using anti-HA beads. Immunoprecipitates and whole-cell extracts were analyzed by immunoblotting with the indicated antibodies.

### PLK1 interacts with CDK1-phosphorylated RAD52 phosphorylated on Thr300

RAD52, a protein which plays a role in MiDAS and double-strand break (DSB) repair (Bhowmick *et al*., 2023; Rossi *et al*., 2021), was present in PLK1-3HA precipitates from mitotic cells (Figs. 1C, 3A, 3D). Interestingly, while investigating RAD52 function in mitosis independently, we generated cells that stably express tetracycline-inducible GFP-tagged RAD52. Cells were synchronized either in G1 (palbociclib) (Fry *et al*, 2004) or prometaphase (nocodazole), and we analyzed GFP-RAD52 precipitates by mass spectrometry to identify interacting proteins (Fig. S5A, B). In this experiment, PLK1 emerged as the top mitosis-specific RAD52-interacting protein (Figs. 6A, S5C). Western blotting of FLAG-RAD52 precipitates confirmed that PLK1 co-immunoprecipitates with RAD52 from cells arrested in mitosis by nocodazole or STLC, while the interaction was minimal in asynchronous cells (Fig. 6B). We also observed endogenous RAD52 interacting with PLK1 in nocodazole-arrested cells (Fig. S5D).

**Figure 6.**
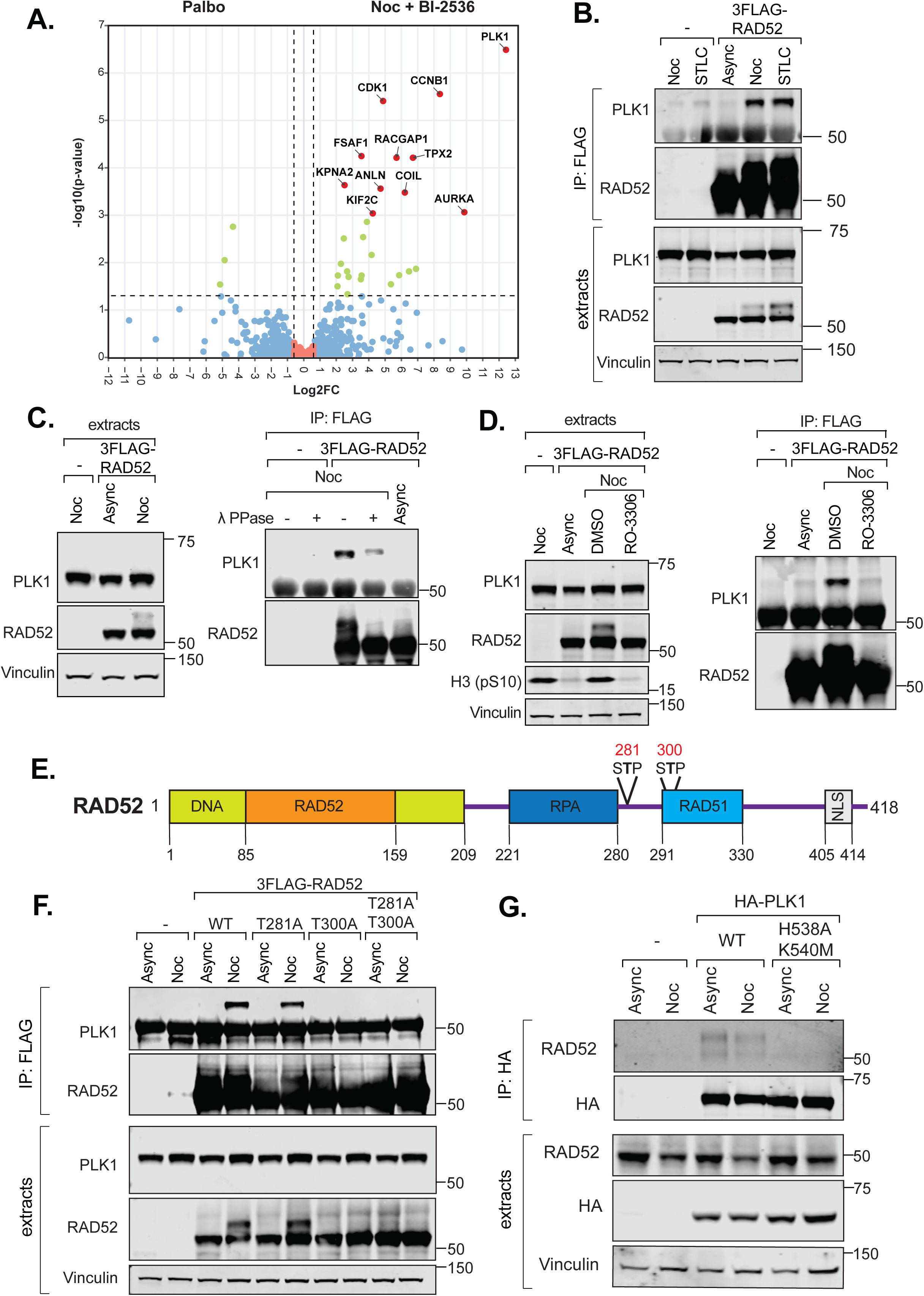
PLK1 interacts with CDK1-phosphorylated RAD52 phosphorylated on Thr300. **A.** Volcano plot of mass spectrometry analysis of GFP-RAD52 immunoprecipitates from U-2 OS FRT cells from 5 biological replicates, showing proteins differentially enriched in G1 cells (left) or G2/M cells treated with BI-2536 (right). The horizontal cutoff line represents a P-value of 0.05. Vertical lines correspond to a fold-change of (±) 1.5. **B.** Extracts from HEK-293 FRT parental cells or cells expressing 3FLAG-RAD52 treated as described were subjected to FLAG immunoprecipitation and immunoblotted for the indicated proteins. Async = asynchronous, Noc = nocodazole. **C.** FLAG immunoprecipitates were either mock-treated or treated with lambda phosphatase, followed by immunoblotting for the indicated proteins. **D.** Nocodazole-arrested HEK-293 FRT 3FLAG-RAD52 cells were treated with DMSO or RO-3306 (10 μM, 15 min). FLAG immunoprecipitates from parental cells treated with nocodazole and from 3FLAG-RAD52 cells treated with DMSO were used as negative controls. The immunoprecipitates were immunoblotted for the indicated proteins. H3 (pSer10) was used as a marker of CDK1 activity. **E.** Schematic diagram showing the domain architecture of human RAD52 and the location of 2 potential polo box binding motifs. **F.** FLAG immunoprecipitates from HEK293 FRT parental and 3FLAG-RAD52 (WT and mutants) cells treated with DMSO or nocodazole were immunoblotted for the indicated proteins. **G.** Extracts from HEK293 FRT parental and HA-PLK1 (WT and H538A K540M) cells treated with DMSO or nocodazole were subjected to HA immunoprecipitation, followed by immunoblotting for the indicated proteins.

As with SLX4, we hypothesised that the PLK1 PBD recognises a CDK1-phosphorylated form of RAD52 present only in mitosis. Supporting this, incubation of FLAG-RAD52 precipitated from mitotic cells with lambda phosphatase greatly reduced the amount of co-precipitating PLK1 (Fig. 6C). Likewise, a short pulse of RO3306 in mitotic cells prevented PLK1 co-precipitation with RAD52 (Fig. 6D). These results indicate a phosphorylation- and CDK1-dependent association between RAD52 and PLK1 in mitosis. We consistently observed a mitosis-specific RAD52 electrophoretic mobility shift, which was reversed by RO3306 and PLK1 inhibitors (Figs. 3A & 6D, S3), or by phosphatase treatment of RAD52 precipitates (Fig. 6C). Thus, the shift in RAD52 during mitosis is attributable to PLK1-dependent phosphorylation.

Human RAD52 contains two potential PLK1 docking motifs, centred on Ser281 and Thr300 (Fig. 6E). As shown in Fig. 6F, a Thr300 to alanine substitution (T300A) abolished co-precipitation of PLK1 with FLAG-RAD52 from nocodazole-arrested cell extracts, whereas substitution of Ser281 had little effect. These findings establish RAD52 as a new PLK1 target in mitotic cells. The T300A mutation also collapsed the RAD52 electrophoretic mobility shift seen in mitosis, indicating that docking of PLK1 is required for PLK1-dependent phosphorylation of RAD52 (Fig. 6F). Furthermore, combined mutation of His538 and Lys540 in the PLK1 PBD severely weakened interaction with RAD52 (Fig. 6G). Overall, these data indicate that RAD52 phosphorylated on Thr300 by CDK1 is a mitotic target of PLK1.

### PLK1-dependent regulation of SLX4 and RAD52 is essential for MiDAS

To probe the functional consequences of PLK1 regulation of SLX4 and RAD52, we focused on SLX4 in U-2 OS cells engineered to express BromoTag-SLX4 (Fig. S2). Degradation of SLX4 by AGB1 led to phenotypic defects that mirrored those observed in previous studies using siRNA depletion of SLX4, or in cells from patients with Fanconi anemia carrying *SLX4* (*FANCP*) mutations (Kim *et al*, 2011; Munoz *et al*., 2009; Stoepker *et al*, 2011). For instance, SLX4 degradation in two independent BromoTag clones resulted in G2 arrest after exposure to the DNA crosslinking agent mitomycin-C (MMC), as well as pronounced hypersensitivity to MMC or cisplatin (Fig. S6), consistent with impaired repair of DNA interstrand crosslinks (ICLs).

To further investigate the effects of SLX4 mutations, we introduced either wild-type SLX4 with an N-terminal GFP tag or a GFP-SLX4 Ser1453Ala mutant (referred to as SLX4) into BromoTag-SLX4 cells via lentiviral transduction. The SLX4 mutant was confirmed as severely defective in its interaction with PLK1 (Fig. 7A). As controls, we used an SLX4 Leu530Ala Trp531Ala double mutant (SLX4), which we previously found to disrupt XPF-ERCC1 interaction and impair ICL repair (Alvarez-Quilon *et al*, 2020), and an SLX4 Cys1805Arg mutant (SLX4), which we previously found to disrupt SLX1 interaction (Castor *et al*, 2013; Nair *et al*, 2014). The expression levels of all SLX4 variants were similar to wild-type, and loss of interaction of each mutant with the relevant partner protein was confirmed (Fig. 7A). As shown in Fig. 7B, degradation of BromoTag-SLX4 led to G2 accumulation after MMC treatment. The GFP-SLX4 mutant could not rescue this defect, while SLX4 restored cell cycle progression to the same extent as GFP-SLX4, as did SLX4. Similarly, SLX4 failed to rescue the pronounced MMC hypersensitivity caused by BromoTag-SLX4 degradation, whereas SLX4 (and SLX4) was similar to wild-type (Fig. 7C). These results indicate that PLK1 docking to SLX4 during mitosis does not influence the role of SLX4 in ICL repair.

**Figure 7.**
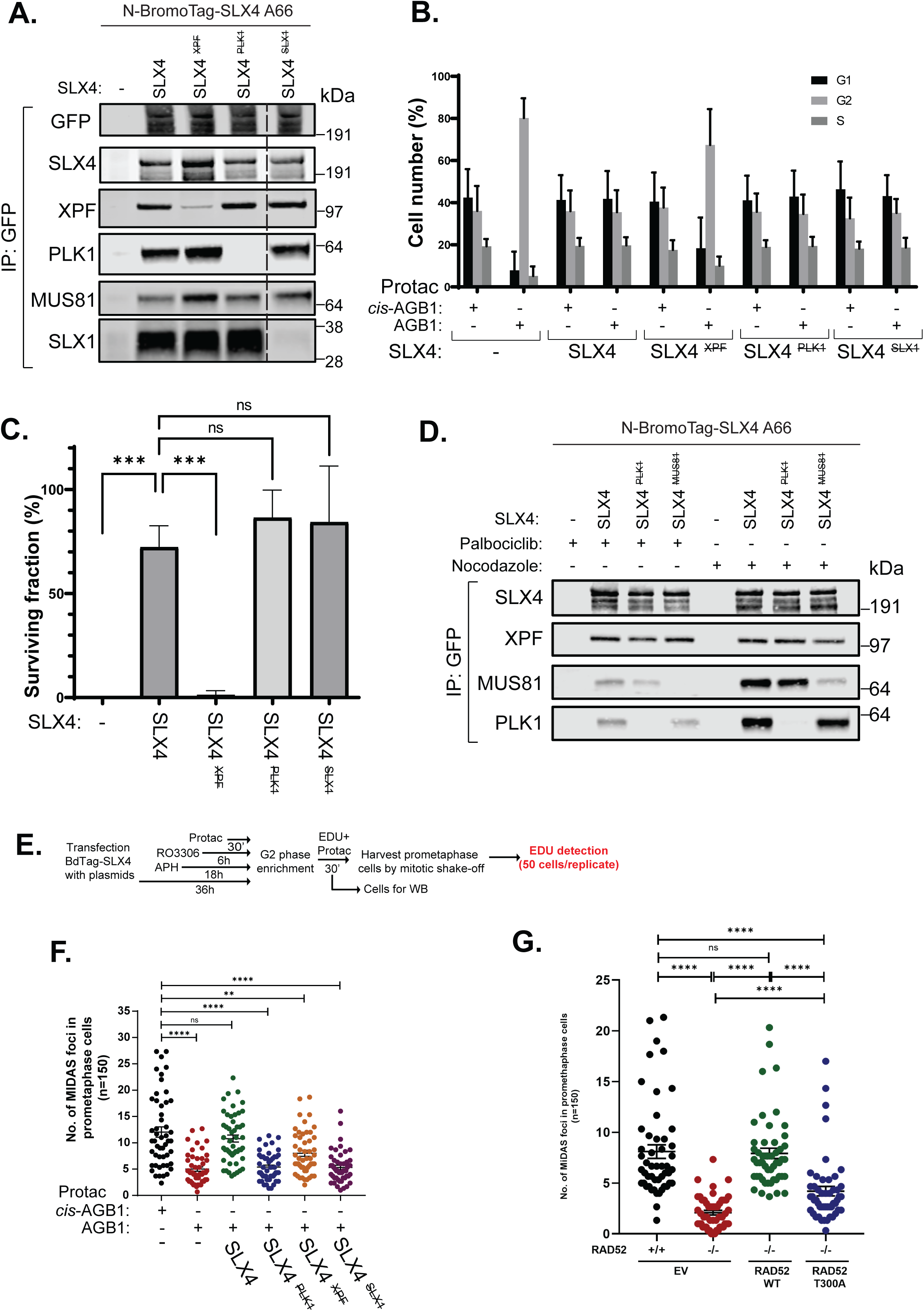
PLK1-SLX4 and PLK1-RAD52 interactions are critical for MiDAS. **A.** SLX4 was immunoprecipitated from extracts of BromoTag–SLX4 cells stably expressing wild-type GFP-SLX4 or GFP-SLX4 mutants defective in binding to XPF, PLK1 or SLX1 using GFP-Trap Sepharose beads. Immunoprecipitates were resolved by SDS-PAGE and analyzed by immunoblotting with the indicated antibodies. An unrelated sample lane was removed from the original blot; the discontinuous line indicates the cropped area. **B.** Stable cell lines shown in (A) were preincubated with either AGB1 or cis-AGB1 (300 nM, 24 h) prior to treatment with MMC (5 ng/mL, 48 h). Cells were then harvested, fixed in ice-cold ethanol, and stained with propidium iodide. Cell cycle distribution was analyzed by flow cytometry. Quantification represents the mean ± SD of three independent experiments. **C.** Surviving fraction of cells following treatment with mitomycin C (10 ng/mL). Stable cell lines shown in (A) were preincubated with either AGB1 or cis-AGB1 (300 nM, 24 h) prior to MMC treatment (10 ng/mL) for 10–14 days. Colonies were then fixed in methanol, stained, and counted. The surviving fraction was calculated as the percentage of colonies in AGB1-treated plates relative to cis-AGB1-treated plates. Quantification represents the mean ± SD of three independent experiments. Statistical significance was determined by one-way ANOVA with Dunnett’s multiple comparisons test. **D.** BromoTag–SLX4 cells stably expressing wild-type GFP-SLX4 or GFP-SLX4 mutants defective in binding to MUS81 or PLK1 were treated with either palbociclib (1 µM) or nocodazole (150 ng/mL). Cell extracts were harvested 24 h later and subjected to immunoprecipitation using GFP-Trap Sepharose beads. Immunoprecipitates were resolved by SDS-PAGE and analyzed by immunoblotting with the indicated antibodies. **E.** Schematic workflow for determining MIDAS proficiency in U-2 OS cells transiently transfected with plasmids that have mutations blocking interaction of SLX4 with PLK1, SLX1, or XPF. **F.** Quantification of number of MIDAS foci in prometaphase cells following treatment as in (E). In. In each quantification, data are presented as mean ± s.e.m. of at least 3 independent experiments. n=150: 50 cells were analyzed in each condition in each replicate. In total, 150 cells were analyzed for each condition. Statistical p values were calculated with two-tailed non-parametric Mann–Whitney tests and indicated in quantification graphs. **G.** Number of EdU foci per prometaphase cells in U-2 OS parental and *RAD52*^-/-^ cells complemented with GFP (EV; empty vector), GFP-RAD52 (RAD52 WT), or GFP-RAD52 T300A (RAD52 T300A). In each quantification, data are presented as mean ± s.e.m. of at least 3 independent experiments. n=150: 50 cells were analyzed in each condition in each replicate. In total, 150 cells were analyzed for each condition. Statistical p values were calculated with two-tailed non-parametric Mann–Whitney tests and indicated in quantification graphs.

Whereas XPF–ERCC1 and SLX1 associate with SLX4 throughout the cell cycle, MUS81–EME1 interacts with the scaffold specifically during the G2/M phases. This mode of temporal regulation juxtaposes MUS81–EME1 with SLX1 on SLX4 in G2/M cells, enabling the coordinated dual incisions required for Holliday junction resolution (Castor *et al*., 2013; Duda *et al*, 2016; Wyatt *et al*, 2013). Previous studies reported that the interaction between MUS81–EME1 and SLX4 depends on CDK1 and PLK1 activity (Wyatt *et al*., 2013), and we wondered if this dependence required the docking of PLK1 on Ser1453-phosphorylated SLX4. However, the SLX4 mutant co-immunoprecipitated MUS81–EME1 from extracts of nocodazole-arrested cells to a similar extent as wild-type SLX4. In contrast, a previously characterized SLX4 [Glu1577Ala+Leu1578Ala] control point mutant (SLX4) markedly reduced the association of SLX4 with MUS81–EME1 (Fig. 7D)

Given that SLX4 has been implicated in MiDAS (Minocherhomji *et al*., 2015), we next asked whether PLK1 docking is required for this process. Cells were treated with aphidicolin to induce replication stress and thereby increase MiDAS frequency, then arrested in G2 with RO3306. BromoTag-SLX4 was degraded using AGB1 PROTAC, and cells were released into mitosis in the presence of EdU; MiDAS was quantified by counting EdU foci in prometaphase cells (Fig. 7E). As shown in Fig. 7F, SLX4 degradation led to a substantial reduction in MiDAS, which could be fully rescued by transient overexpression of wild-type SLX4. In contrast, the SLX4 mutant was unable to rescue the MiDAS defect, and these cells resembled the vector-only negative controls. These data suggest that the SLX4 mutation represents a separation-of-function allele, specifically disrupting MiDAS without affecting other functions of SLX4. It interesting to note that both the SLX4 and SLX4 mutants showed partial impairment in MiDAS (Fig. 7F).

Similar to SLX4, RAD52 has also been implicated in MiDAS in previous studies (Audrey *et al*, 2024; Bhowmick *et al*, 2016; Minocherhomji *et al*., 2015). This prompted us to ask whether the interaction between PLK1 and RAD52 is functionally important for this pathway. To address this, we generated U-2 OS cell lines lacking RAD52 (Fig. S7) and subsequently re-expressed either wild-type RAD52 or a Thr300Ala (T300A) mutant in the *RAD52*^-/-^background. As shown in Fig. 7G, the frequency of MiDAS events was markedly reduced in RAD52-deficient cells compared to parental controls. Re-expression of wild-type RAD52 restored MiDAS activity, whereas expression of the Thr300Ala mutant provided only partial rescue. These findings indicate that PLK1-mediated regulation of RAD52, as well as SLX4, is critical for efficient MiDAS. Thus, PLK1 emerges as a central regulator of this pathway.

## Discussion

This study demonstrates that Polo-like kinase 1 (PLK1) regulates a broader mitotic genome-maintenance network than previously recognised. Using a substrate-trapping approach with acute PLK1 inhibition, we isolated endogenous (HA-tagged) PLK1 complexes from mitotic cells and identified associated proteins enriched for roles in chromosome segregation, DNA replication, and DNA repair. Several known factors in mitotic genome maintenance, including SLX4, POLQ, and RHINO, were identified as interactors, validating our method. Notably, we also discovered previously uncharacterised mitotic PLK1 interactors relevant to genome stability, including RAD52, REV1, and FANCM. Focusing on SLX4 and RAD52, our data show that both bind PLK1 via canonical CDK1-dependent phospho-motifs that are necessary for effective PLK1 interaction during mitosis. Functional studies further indicated that disrupting PLK1 docking specifically compromises mitotic DNA synthesis (MiDAS); in the case of SLX4, its role in ICL repair appeared to be unaffected by this change. Collectively, these results support a model in which PLK1 functions not only as a mitotic kinase overseeing chromosome segregation but also as a coordinator of mitotic responses to under-replicated or damaged DNA.

Although the precise sites in SLX4 and RAD52 phosphorylated by PLK1 remain unmapped, the available data indicate that their modification is docking-dependent. The loss of mitotic mobility shift in the RAD52 Thr300Ala mutant is particularly telling: the wild-type RAD52 shift is abolished by PLK1 inhibitors and reversed by phosphatase treatment, indicating that it reflects PLK1-catalysed phosphorylation. Together, these findings support a model in which loss of docking prevents PLK1 from effectively engaging RAD52 and other substrates, thereby inhibiting their phosphorylation. This is consistent with the established model in which CDK1-dependent phosphorylation generates a docking site for the PBD, thereby positioning the PLK1 kinase domain to target nearby residues (Elia et al., 2003b; Lowery et al., 2005). Many characterised PLK1 substrates operate similarly, with disruption of docking blocking downstream PLK1-dependent phosphorylation events (Elia et al., 2003b; Preisinger et al., 2005). While future phosphoproteomic analyses will define the exact residues in RAD52 - and SLX4 - modified following docking, our current data establish the critical mechanistic role of this interaction.

Our results further suggest that PLK1-mediated regulation of SLX4 and RAD52 is highly specific to mitotic genome maintenance, rather than required for all functions of these proteins. This specificity is particularly clear for SLX4. The SLX4 Ser1453Ala mutant restored resistance to MMC and prevented G2 arrest after loss of endogenous SLX4, suggesting that recruitment of PLK1 to SLX4 is not necessary for ICL repair. However, this same mutation did not rescue MiDAS, defining a separation-of-function allele that uncouples the mitotic role of SLX4 from its interphase DNA repair activity. A similar pattern was observed for RAD52, where loss of PLK1 docking impaired its ability to facilitate MiDAS. Together, these findings suggest that PLK1-dependent regulation of SLX4 and RAD52 constitutes a specific mitotic control mechanism.

An additional outcome of this work is the generation of a conditional human SLX4 loss-of-function system that should prove broadly useful for the study of SLX4 biology. Although *Slx4*/*Btbd12* knockout mice are viable and appear normal (Castor *et al*., 2013), we and others have repeatedly been unable to recover complete SLX4 knockout clones from a wide range of cells, suggesting that complete loss of SLX4 may be poorly tolerated in human cells (data not shown). By contrast, BromoTag-mediated acute degradation enabled efficient and reversible depletion of endogenous SLX4, allowing rigorous complementation analyses with separation-of-function mutants such as SLX4 Ser1453Ala. This approach overcomes a longstanding experimental limitation in the field and should provide a valuable platform for dissecting the multiple genome maintenance functions of SLX4 in human cells.

A notable implication of our findings is that PLK1 docking to SLX4 is not required for recruitment of MUS81–EME1 to the SLX4 scaffold. Previous work showed that association of MUS81–EME1 with SLX4 is enhanced in G2/M and depends on CDK1 and PLK1 activity, leading to the proposal that PLK1-mediated phosphorylation events promote assembly of the SLX4–MUS81–EME1 complex during mitosis (Wyatt et al., 2013). Because we identified Ser1453 as the principal PLK1 docking site in SLX4, we considered the possibility that PLK1 recognition of CDK1-phosphorylated SLX4 might directly mediate MUS81–EME1 recruitment. However, the SLX4 Ser1453Ala mutant retained normal interaction with MUS81–EME1 in mitotic cells, whereas disruption of the previously characterised MUS81-binding region in SLX4 strongly impaired this association (Fig. 7D). These observations indicate that PLK1 docking to SLX4 is mechanistically distinct from PLK1-controlled MUS81–EME1 recruitment to the scaffold. Instead, PLK1 engagement with SLX4 likely regulates downstream mitotic functions of the complex after assembly has occurred – perhaps PLK1 bound to SLX4 activates MUS81-EME1 once it binds to the scaffold, for example.

How might PLK1 regulation of SLX4 and RAD52 influence the functions of these proteins? One possibility is that PLK1-dependent phosphorylation may modulate the enzymatic activity or substrate specificity of its targets. Uncontrolled nuclease activity during mitosis could be damaging, especially on condensed chromosomes under spindle tension. Likewise, the recombination activities of RAD52 might need to be limited to prevent the formation of abnormal DNA structures during chromosome segregation. Thus, PLK1 phosphorylation could act as a molecular switch that directs these proteins towards mitotic functions compatible with chromosome segregation, while restraining inappropriate repair activities. In this light, it is interesting to note that the SLX4 and SLX4 mutants are similar to the SLX4 mutant in that they reduce MiDAS frequency. This observation raises the possibility that PLK1 somehow controls the activity of XPF-ERCC1 and SLX1 bound to SLX4 in mitosis in a way that is required to facilitate MiDAS.

Our results also identify PLK1 as a potentially central regulator of MiDAS. Previous studies have implicated several of the proteins we identified—including SLX4, MUS81-EME1, RAD52, and POLQ—in MiDAS. Our data suggest that PLK1 may orchestrate multiple components of this pathway simultaneously. Rather than acting through a single downstream effector, PLK1 may serve as an organising kinase that assembles and activates a mitotic DNA-processing network tailored to complete replication under mitotic conditions. The discovery of FANCM and REV1 as mitotic PLK1 interactors further broadens this framework. FANCM is known to remodel stalled replication forks, while REV1 serves as a scaffold for translesion synthesis polymerases. The observation that the PLK1-FANCM interaction is most evident after acute PLK1 inhibition suggests that FANCM may engage in rapid cycles of docking, phosphorylation, and release. REV1 may be regulated in a similar manner. While the specific functional outcomes remain to be clarified, PLK1-dependent regulation of these proteins may facilitate replication fork processing, lesion bypass, and completion of DNA synthesis at challenging genomic regions during mitosis.

More generally, these findings support the emerging view that PLK1 operates at the intersection of mitotic progression and DNA damage responses. While PLK1 has long been regarded as a cell-cycle kinase responsible for processes such as centrosome maturation, kinetochore function, and cytokinesis, recent evidence shows that it also plays crucial roles in checkpoint recovery and DNA repair pathway selection. Our data extend this concept by identifying multiple DNA repair and replication-associated proteins as mitotic PLK1 targets and by establishing the functional significance of PLK1 regulation in MiDAS. Several key questions remain. The exact PLK1-dependent phosphorylation sites in SLX4 and RAD52 have not yet been determined, and it will be important to assess how these modifications affect localisation, complex assembly, and biochemical function. It is also necessary to clarify whether PLK1 directly phosphorylates FANCM, REV1, and other candidate proteins identified in our proteomic studies. Given that cancer cells rely on tolerance to replication stress, elucidating how PLK1 coordinates these pathways could have therapeutic relevance. PLK1 inhibitors are under clinical investigation, and our findings raise the possibility that their effectiveness may, in part, result from disrupting mitotic DNA synthesis and the processing of under-replicated DNA.

## Ackowledgements

We thank the excellent technical support of the MRC PPU including the Mass Spectrometry Team, the DNA Sequencing Service, Tissue Culture Team, and the Reagents and Services Team. We thank Arlene Rennie and Rosemary Clarke (Centre for Advanced Scientific Technologies, School of Life Sciences Dundee) for extensive help with FACS-based cell sorting and flow cytometry experiments. We thank all JR lab members for useful discussions. This work was supported by the Medical Research Council (grant number MC_UU_12016/1; J.R.) and by a PhD studentship funded by Merck KGaA (D.U.), European Union Marie Skłodowska-Curie Actions grant (Antihelix 859853, Y.L.), Danish Independent Research Fund (1030-00180B; Y.L.) and NEYE Foundation (122479, Y.L).

## Author contributions

D-C.U. and I.M. did the majority of the experiments in this study. W.B. made the SLX4 degron cell lines. T.M. made the genome editing constructs and helped with genotyping. S.A.B. and Y.L. were responsible for the MiDAS analyses. Y.W. made the PLK1-3HA cell line. F.L. carried out the mass spectrometric applications, and F.W. analysed the mass spectrometry data and wrote the relevant scripts. J.R. conceived the project with input from I.M, D-C.U. and W.B. and also wrote the paper.

## Materials and Methods

### Reagents

All reagents including cDNA clones, antibodies, peptides and oligonucleotides mentioned in the present study can be found in Table S2.

### Cell lines and drug treatments

U-2 OS, HCT116, and HEK-293 cells were grown in Dulbecco’s modified Eagle’s medium (DMEM) (Sigma-Aldrich) supplemented with 10% foetal bovine serum (FBS) (Sigma-Aldrich), 1% penicillin/streptomycin 10,000 U/ml (Gibco), 1% L-glutamine 200 mM (Gibco), 1 mM sodium pyruvate (Gibco), and 1X non-essential amino acids (Gibco). U-2 OS Flp-In T-REx and HEK-293 Flp-In T-REx cells with integrated transgenes were grown in the same media as above supplemented with hygromycin (200 μg/ml) and blasticidin (15 μg/ml).

To synchronize cells in G1, palbociclib (1 μM) was added for 16h (HCT116), 18h (HEK-293) or 20h (U-2 OS). To synchronize cells in G2/M, nocodazole (150 ng/ml) was added for 16h (HCT116), 18h (HEK-293) or 20h (U-2 OS). S-trityl-L-cysteine (STLC) was added at a final concentration of 5 μM for 20h. To inhibit PLK1 activity in mitosis, nocodazole-arrested cells were treated with BI-2536 (500 nM) or BI-6727 (500 nM) for 1 hour. To inhibit CDK1 activity in mitosis, nocodazole-arrested cells were treated with RO-3306 (10 μM) for 15 minutes or Flavopiridol (10 μM) for 15 minutes.

To degrade target protein BromoTag® AGB1 (Biotechne, Cat. 7686; 300-500nM), BromoTag® cis-AGB1 (Biotechne, Cat. 7687; 300-500nM) were added to the cells prior to experiment.

### Inducible protein expression in Flp-In T-REx cells

Stable protein expression in U-2 OS Flp-In T-REx and HEK-293 Flp-In T-REx cells was achieved by co-transfection of pcDNA5/FRT/TO plasmid (1 μg) and POG44 (9 µg) with PEI (Polysciences), according to the manufacturer’s instructions. The cells that integrated the FRT plasmid were selected with hygromycin (200 μg/ml) and blasticidin (15 μg/ml). The expression of the desired protein was induced by treatment with either tetracycline or doxycycline as indicated.

### CRISPR/Cas9 genome editing

HCT116 and U-2 OS cells (seeded at 50%-60% confluence in 10cm dishes) were transfected using commercially available PEI (Polysciences), according to the manufacturer’s instructions. To generate *RAD52^-/-^* cells, 1 μg of plasmid encoding sense gRNA (DU 74245) and 1 μg of plasmid encoding antisense gRNA and Cas9 nickase (DU 74246) were used. For engineering *PLK1-3HA* cells, 2 μg of plasmid encoding a gRNA and wild-type Cas9 (DU 81937) and 3 μg of plasmid encoding the PLK1-3HA donor template (DU 81400) were used. To generate BromoTag-SLX4 cells, 1.5 µg pX459 plasmid encoding the single guide RNA and wild-type Cas9 (DU74465) and 3 µg pMK plasmid encoding the donor sequence (DU74456) were used. A transfection mixture was prepared by combining plasmid DNA with PEI and OPTIMEM Reduced Serum Medium (ThermoFisher, Cat. 31985070). The transfection mixture was briefly vortexed, followed by a 15 min incubation at room temperature.

During the incubation, the cell culture media was replaced with antibiotics-free media. The transfection mixture was added dropwise to the culture media, and the cells were placed back into the incubator. 24h later, the cell culture media was changed to normal media supplemented with 2 μg/ml puromycin. The puromycin media was refreshed after 24 h for a total selection time of 48h. The cells were then allowed to recover for a few days, and then single cells were isolated by fluorescence-activated cell sorting (FACS). In the case of *RAD52^-/-^* cells, single cells were isolated, whereas GFP-positive single cells were isolated for PLK1-3HA and BromoTag-SLX4 cells. The single cell clones were grown until sufficient material was obtained for Western blotting or genotyping by PCR. Due to percentage of GFP-positive cells for BromoTag-SLX4, two rounds of transfection were performed, then the transfected cell pools were bulk sorted GFP-positive cells using the standard semi-purity mode, expanded, and subsequently single cell sorted as described above.

### Genotyping of clones

For genotyping of BromoTag-SLX4 clones, the *SLX4* locus was amplified using the Terra Direct PCR Red Dye Premix kit (Takara Bio) following the manufacturer’s instructions using primers SLX4 Nter Fwd + SLX4 Nter Rev. When clones reached 50-80% confluence in the 96-well plate, cells were harvested by trypsinisation. Half of the cells were collected in a sterile microcentrifuge tube for screening by PCR, while the cells were seeded in a 6-well plate and expanded for further screening by western blot. The cells in the microcentrifuge tube were collected by centrifuging at 500g for 5 minutes, the supernatant was aspirated carefully, then the cell pellet was resuspended in 20 μL PBS and used as a template for PCRs.

For genotyping of *RAD52^-/-^* clones, genomic DNA was extracted from cells using the Blood and Tissue DNeasy kit (Qiagen), according to the manufacturer’s instructions. The *RAD52* locus was amplified by PCR using the Q5 High-Fidelity DNA Polymerase (NEB) was used, according to the manufacturer’s instructions, with primers ex4 F and ex4 R. PCR products were loaded on a 1% (w/v) agarose gel in Tris–acetate-EDTA (TAE) buffer (40 mM Tris, 20 mM acetic acid, 1 mM EDTA, pH 8.0) with 1X SYBR™ Safe DNA Gel Stain (Thermo Fisher). 5-10 μl of 1 kb Plus DNA Ladder (Thermo Fisher) was used as a marker for size of DNA samples. DNA samples were separated through electrophoresis in TAE buffer at 100V for 45 min and gels were imaged using U:Genius Gel documentation system (Syngene) or ChemiDoc Imaging System (BioRAD) and analysed in Image Studio.

When amplifying the junctions of the targeted *SLX4* locus, the following primers were used: SLX4 Nter Fwd + GFP Rev to amplify the 5’ junction, BromoTag Fwd + SLX4 Nter Rev to amplify the 3’ junction, or SLX4 Nter Fwd + SLX4 Nter Rev to amplify the *SLX4* target knock-in locus. The elongation time was limited to selectively amplify the wild-type allele but not the knock-in allele with the latter primer set.

The amplicon was cloned into the pSCB plasmids using the StrataClone PCR cloning Kit (Agilent) following the manufacturer’s instructions. Approximately 15 bacterial colonies were picked for each single cell clone, and overnight bacterial cultures were prepared in 5 mL LB media supplemented with 100 μg/ml ampicillin at 37 °C, shaking. Bacterial cultures were pelleted at 5000 rpm for 15 min and plasmid DNA was isolated by DNA Sequencing and Services (University of Dundee). Plasmids with correct insertion of the PCR product, identified by restriction enzyme digestion of the plasmid with *EcoRI*-HF (New England Biolabs) according to manufacturer’s instructions, were sequenced with M13 forward and reverse primers. The sequencing was carried out in-house by the MRC PPU DNA Sequencing and Services (MRC Reagents & Services) using Applied Biosystems automated DNA sequencers. Data was analysed using SnapGene Viewer (v5.3.0).

### Cell lysis and immunoprecipitation

For protein extraction, cells were lysed at 4°C for 30 min in Mammalian Lysis Buffer containing 50 mM Tris-HCl pH 7.5, 150 mM NaCl, 1% (v/v) Triton-X (Sigma-Aldrich), 0.5% (v/v) NP-40 alternative (Calbiochem), 270 mM sucrose, 1:5000 Pierce Universal 20 Nuclease (ThermoFisher Scientific), Phosphatase inhibitor (1:100 Phosphatase Inhibitor Cocktail 2 (Sigma-Aldrich) or PhosStop (Roche), 1x Complete EDTA-free protease inhibitor (Roche), 1 mM DTT or 1% β-mercaptoethanol, and 10 ng/ml Microcystin-LR (Sigma-Aldrich). The mixture was then centrifuged at 17000g, 4°C for 15 min and total protein concentration was quantified using Bradford assay.

To immunoprecipitate epitope-tagged proteins, HA-Trap magnetic agarose (Proteintech, atma), GFP-Trap magnetic agarose (Proteintech, gtma), or FLAG M2 agarose beads (Sigma-Aldrich, A2220) were washed three times with lysis buffer and once in PBS, then incubated with protein lysates for 2-3h at 4°C with rotation. To immunoprecipitate RAD52 or PLK1, 10 µL protein A magnetic dynabeads (ThermoFisher Scientific, 10001D) were washed twice with PBS containing 0.02% (v/v) Tween-20 (Sigma-Aldrich). The beads were then incubated with anti-PLK1, anti-RAD52 antibody or rabbit isotype control for >30 min at room temperature with agitation. The antibody-conjugated beads were then incubated with cell extracts for 2-3 hours at 4°C with agitation. To immunoprecipitate SLX4, 20 µL protein A/G agarose bead slurry (50% in PBS; MRC Reagents & Services) was incubated with anti-SLX4 antibody or sheep isotype control for >30 min at room temperature with agitation. Clarified cell extracts were pre-cleared by diluting to 2 mg/mL in Mammalian Lysis Buffer, followed by incubation with 20 µL protein A/G agarose beads for 15-30 minutes at 4°C with agitation. Subsequently, samples were centrifuged at 2000g 4°C for 1 min, and the pre-cleared cell extracts (supernatant) were transferred to new 1.5 mL microcentrifuge tube along with the antibody-coupled beads and incubated for 1.5-3h at 4°C with agitation. After incubation, the beads were washed 3-4 times with ice-cold IP Wash Buffer (50 mM Tris-HCl pH 7.4, 250 mM NaCl, 270 mM sucrose, 1% (v/v) Triton X-100 (Sigma-Aldrich), 0.5% (v/v) NP-40 alternative (Calbiochem), then eluted as above and stored at –20°C until western blotting.

### Western blotting

Protein lysates were mixed with LDS sample buffer (Invitrogen) and incubated at 95°C for 5-10 min. 20-50 µg of total protein were loaded on each lane of a NuPAGE^TM^ 4-12%, or 8%, or 12% Bis/Tris gel (ThermoFisher Scientific), or a NuPAGE^TM^ 3-8% Tris-acetate gel (ThermoFisher Scientific) and run at 150V for 1-1.5hrs. The proteins were then transferred onto a 0.45 µm nitrocellulose membrane (Amersham) at 90V for 90 min. The membranes were then blocked in 5% milk in TBS-Tween (0.1% v/v) and then incubated with primary antibodies overnight at 4°C. This was followed by 3×5 min washes with TBS-Tween (0.1% v/v) and incubation with secondary antibodies for 1h at room temperature. The membranes were then washed with TBS-Tween (0.1% v/v) (3×5 min), and imaged. When antibodies recognizing phosphorylated antigens were used, 3% BSA (Sigma-Aldrich) in TBS-Tween (0.1% v/v) was used for blocking and antibody incubations. When a light chain-specific secondary antibody was used for western blotting, blocking and antibody incubations were performed using 5% normal mouse serum (ThermoFisher Scientific) in TBS-Tween (0.1% v/v).

For immunoblotting with the phospho-specific SLX4 pSer1453 antibody, the antibody was diluted to a final concentration of 0.5 μg/mL in 5% (w/v) non-fat dry milk in TBS-Tween (0.2% v/v) plus 40 μg/mL of the corresponding non-phosphorylated peptide. The antibody-peptide solution was incubated for 1h at room temperature prior to use.

### Mass spectrometry

To prepare samples for mass spectrometry, proteins were bound to S-Trap columns (Protifi) and processed on the column according to the manufacturer’s instructions, with the following modifications. After eluting the proteins from the beads with 5% SDS, the reduction and alkylation steps were performed within a single step. 2 µl TCEP 125 mM (final concentration: 5 mM) and 2 µl 2-chloroacetamide 500 mM (final concentration: 20 mM) were added and the samples were then incubated at 37°C for 10 min with shaking. For protein digestion, Digestion buffer was prepared by reconstituting 100 µg of lyophilised Trypsin/LysC (ThermoFisher Scientific) into 50 mM acetic acid and further diluted to 0.05 µg/µl in TEAB 100 mM. 20 µl of digestion buffer was added to each sample, and the columns were closed loosely to reduce evaporation but not create airtight seal. The proteins on the columns were digested overnight at 37°C. Next day, the peptides were eluted off the columns with 40 µl TEAB 100 mM, followed by 40 µl formic acid 0.2%, and subsequently with 40 µl ACN 5%. The lyophilised peptides were resuspended in 5% formic acid in water, then injected on an UltiMate 3000 RSLCnano System coupled to an Orbitrap Fusion Lumos Tribrid Mass Spectrometer (ThermoFisher Scientific). Samples were loaded on an Acclaim Pepmap trap column (ThermoFisher Scientific #164750) prior analysis on a PepMap RSLC C18 analytical column (ThermoFisher Scientific #ES903) and eluted with a 120 min linear gradient from 3 to 35% of buffer comprising of 0.08% formic acid in 80:20 (v:v) acetonitrile in water. Subsequently, eluted peptides were analysed by the mass spectrometer operating in data independent analysis mode.

### PLK1-3HA IP/MS data analysis

Results from the DIA-NN search were processed using R (version 4.5.1) (Team, 2023) in the RStudio environment (version 2025.09.2+418) (Posit, 2024) under application of in-house written scripts, with some parts based on scripts used in (Khanam *et al*, 2021). In brief, protein intensity data was transformed and calibrated using variance stabilizing normalization (VSN) (Huber *et al*, 2003; Huber *et al*, 2002) (Figure S8). Proteins were statistically tested using limma under application of robust hyperparameter estimation (Phipson *et al*, 2016; Ritchie *et al*, 2015) and filtered using an adjusted p-value cut-off of 0.05 (5% false discovery rate (FDR)). In case the moderated t-statistic could not be calculated due to insufficient (zero) observations in the negative control group (HA immunoprecipitation from untagged parental cells), a protein was classified “total-regulation” if it was detected in at least 4 out of 6 PLK1-3HA replicates and for such observations the intensity of the control samples was set to 1 (log_2_ = 0) to be able to estimate a log_2_ fold change for result ranking. Gene Ontology (GO) term analysis was performed using the clusterProfiler library (Xu *et al*, 2024) and terms were combined based on their semantic similarity (Wang *et al*, 2007a; Yu *et al*., 2010), using a cut-off value of 0.65. Protein-interaction network analysis was conducted using the STRINGdb library (Szklarczyk *et al*, 2023). For this, protein interactions exhibiting a combined score ≥ 700 (“high confidence”) were retained and filtered for proteins possessing either of the following GO terms: GO:0006974 (DNA damage response), GO:0006281 (DNA damage repair), GO:0006260 (DNA replication), further for GO:0006974 also the offspring terms were included. All R scripts used for data analysis can be downloaded from Zenodo (European Organization For Nuclear & OpenAire, 2013) under https://doi.org/10.5281/zenodo.20156926, while the mass spectrometry raw data can be accessed via PRIDE under the accession number PXD080479 (Perez-Riverol *et al,* 2025). Data used for statistical testing and associated GO terms can be accessed and explored via CURTAIN (Phung *et al*, 2024) under https://curtain.proteo.info/#/42c957a8-184e-4fb0-b9a7-80f1bafc0687. Following R libraries were used: stringr (Wickham, 2023), dplyr (Wickham *et al*, 2023a), vsn, limma, ggplot2, reshape2 (Chang & Bertrand, 2025; Wickham, 2007), viridisLite (Garnier *et al*, 2023), ggrepel (Slowikowski, 2024), ggpointdensity (Kremer, 2019), scales (Wickham *et al*, 2023b), ggnetwork (Briatte, 2024), STRINGdb, AnnotationHub (Morgan & Shepherd, 2024), clusterProfiler and GO.db (Carlson, 2025).

### GFP-RAD52 IP/MS data analysis

The spectral library used for peptide identification was in-house predicted by DIA-NN (version 2.3.2) (Demichev *et al*, 2020; Steger *et al*, 2021) from a *Homo sapiens* Uniprot FASTA (42,583 entries (including isoforms), downloaded 29/01/2026 from www.uniprot.org), with additional inclusion of contaminants (as per DIA-NN). For the spectral library prediction, the protease was set as trypsin under omission of the proline rule, with a maximum of 2 missed cleavages, setting of carbamidomethylation of cysteine as fixed modification, and a maximum 3 of the following variable modifications: Protein N-terminal methionine excision, protein N-terminal acetylation, phosphorylation (STY) and oxidation of methionine. Peptide length was set to 7-30 residues, precursor charge 2-5, precursor m/z range from 380-980 and fragment ion range was set to 150-2000 m/z ratio. Scoring was set as proteoforms and the proteotypicity set as isoform IDs with an FDR of 1%. The raw mass spectrometry data was searched using DIA-NN (version 2.3.2) using the same settings as for spectral library prediction, however using a global 5% FDR. Results were further processed using R (version 4.5.1) using in-house written scripts, with some parts based on (Khanam *et al*., 2021). In brief, peptides intensities were averaged (in case of multiple detections) and the protein intensity was calculated as the sum of all individual peptide intensities. Afterwards, the intensities were VSN transformed and the individual samples calibrated to each other (Figure S9). Statistical analysis was conducted using limma with robust hyperparameter estimation and a 5% FDR. Background proteins were subtracted (either higher abundant in GFP-only control or not differentially abundant to the GFP-RAD52 samples). All other analyses (e.g. “total regulations”) were conducted as described for the PLK1-3HA IP/MS. All mass spectrometry raw data, spectral library, search engine results can be downloaded from ProteomeXchange or jPOSTrepo via the accession numbers JPST004626 and PXD078271. Data analysis scripts are available from Zenodo under the URL https://doi.org/10.5281/zenodo.20156926, while the results can be interactively explored via CURTAIN under https://curtain.proteo.info/#/fd26cd40-d713-4f8b-9524-e12130e11ba3.

### Cell cycle analysis by flow cytometry

Cells were harvested by trypsinization, centrifuged, and washed with PBS. For nocodazole-and STLC-treated cells, floating cells in the cell culture media were also collected. The cell pellets were fixed with ice-cold 70% ethanol for a minimum of 15 minutes at −20°C. DNA was stained using propidium iodide (PI, ThermoFisher Scientific; 50 μg/mL) and RNase A (ThermoFisher Scientific; 100 µg/mL) or DAPI (Invitrogen; 5 μg/mL) for 30 minutes at room temperature in the dark. The samples were analysed using the BD LSRFortessa or BD FACSCanto^TM^ II flow cytometer. Data analysis was performed using the FlowJo software (v. 10.10.0).

### Clonogenic survival assay

Cells were seeded at 1000 cells per well in 6-well plates and allowed to attach for 4-6 hours. Subsequently, cells were treated with 300 nM AGB1 or cis-AGB1 for 18–24 h, followed by treatment with 0–20 ng/mL mitomycin C (MMC; Sigma) for 24 h. For genotoxin sensitivity assays using BromoTag–SLX4 cell lines rescued with GFP-SLX4 point mutants, transgene expression was induced by addition of tetracycline at a final concentration of 10–30 ng/mL concomitantly with PROTAC treatment. After 24 hours, the culture media was replaced with genotoxin-free complete DMEM. Cells were then incubated for 7-10 days until visible colonies had formed. Plates were washed twice with 1xPBS, air-dried, fixed with 100% methanol, stained with crystal violet, then washed with water. The number of colonies was manually counted. Data analysis was performed using Graphpad Prism 10. Three technical replicates were made per condition, and a minimum of three independent experiments were performed.

### Analysis of G_2_/M-arrest

Cells were seeded at 1.25×10^5^ cells per well in 6-well plates and allowed to adhere for 4-16 hours. Next, cells were treated with 300 nM AGB1, 300 nM *cis*-AGB1, or DMSO. After 6-16 hours, MMC was added to the cells, which were further incubated for another 48 hours. Cells were harvested with trypsin, washed with PBS, then fixed with ice-cold 70% (v/v) ethanol for 30 minutes at 4°C, and stored at 4°C until processing. Cell cycle profiles were examined as outlined above.

### Characterisation of BromoTag-SLX4 clones

Cells were seeded in 10 cm dishes at low confluence and allowed to attach overnight, then treated as indicated below.

For the AGB1 dose-response, cells were treated for 3 hours with AGB1 at a final concentration of 0, 1, 3, 10, 30, 100, 300, or 1000 nM, *cis*-AGB1 at a final concentration of 1000 nM, or DMSO. For the AGB1 time-course, cells were treated with 300 nM AGB1 for 0, 1, 3, 6, or 24 hours, or for 24 hours with 300 nM *cis*-AGB1 or DMSO. To assess dependence on NEDDylation and proteasome, cells were pre-treated with 3 μM of MLN4924 (Active Biochem), 50 μM of MG132 (Sigma), or DMSO for 1 hour. Subsequently, AGB1 or *cis*-AGB1 was added to a final concentration of 300 nM, or equal volume of DMSO, then cells were incubated for an additional 3 hours. After treatment, cells were washed twice with PBS, then plates were stored at −20°C. Cells were lysed, subject to immunoprecipitation for SLX4, followed by western blotting.

### Plasmid transfection

Depending on the experiment, cells were transfected using either PEI or FuGENE according to the manufacturer’s instructions. HEK293 cells were transfected using the HBS/CaCl₂ method as previously described (Munoz *et al*, 2018).

### Mitotic DNA synthesis (MiDAS) detection

To detect MiDAS, we followed a published protocol (Garribba *et al*, 2018). Briefly, culture cells were first treated with 0.3 μM APH and arrested at G2 phase with CDK1 specific inhibitor RO3306 (APExBIO; 7µM) and then released into mitosis for an enrichment of mitotic cells. The duration of the treatment is shown in the flowchart of the experiment. Cells were then washed with pre-warmed (37°C) PBS (ThermoFisherScientific) three times within 5 minutes and subsequently released into pre-warmed (37°C) culture medium containing EdU (ThermoFisher Scientific; 40 µM) for 30minutes. The ‘round-up’ mitotic cells were then harvested by mitotic shake-off and subjected to EdU detection by Click-IT chemistry (Click-IT EdU; Alexa fluor 594, Imaging Kits, Life Technologies). Slides were mounted with DAPI-containing Vectashield (Vector Laboratories). Images with EdU foci were captured using an Olympus BX63 microscope and analyzed with CellSens Package (Olympus), and MiDAS foci were counted manually with samples unknown to the counter.

### Peptide pull-downs

For peptide pull-down assays, biotinylated peptides were incubated with streptavidin beads for 2 h at 4°C at a ratio of 5 µg peptide per 10 µL beads in peptide buffer (50 mM Tris-HCl, pH 7.9, 100 mM NaCl). Beads were washed three times with the same buffer prior to use in subsequent experiments.

Recombinant 6His-PLK1 was precleared by incubation with empty streptavidin beads at a 1:1 (v/v) ratio. For each pull-down, 20 µg of recombinant PLK1 was incubated with peptide-coupled beads for 2 h at 4°C. Beads were then washed five times with lysis buffer, followed by a final wash in PBS, and resuspended in LDS sample buffer.

For pull-downs using HEK293 cell lysates, 100 µg of extract was incubated with 10 µL of peptide-coupled beads for 2 h at 4°C. Beads were subsequently washed and processed as described above.

### Lambda phosphatase treatments

Lambda-Phosphatase treatment was performed using the Lambda Protein Phosphatase kit (New England Biolabs) according to the manufacturer’s instructions. Briefly, immunoprecipitates were washed three times with wash buffer and once with λ-phosphatase buffer prior to incubation with lambda-phosphatase (400 U) for 30 min at 30°C with gentle shaking. For phosphatase inactivation, lambda-phosphatase was preincubated with EDTA (100 mM) and microcystin-LR (10 ng/mL) for 30 min at 4°C prior to use.

